# The Role of Action Potential Waveform in Failure of Excitation Contraction Coupling

**DOI:** 10.1101/2021.06.29.450361

**Authors:** Xueyong Wang, Murad Nawaz, Steve R.A. Burke, Roger Bannister, Brent D. Foy, Andrew A. Voss, Mark M. Rich

## Abstract

Excitation contraction coupling (ECC) is the process by which electrical excitation of muscle is converted into force generation. Depolarization of skeletal muscle resting potential contributes to failure of ECC in diseases such as periodic paralysis, ICU acquired weakness and possibly fatigue of muscle during vigorous exercise. When extracellular K^+^ is raised to depolarize the resting potential, failure of ECC occurs suddenly, over a range of several mV of resting potential. While some studies have hypothesized the sudden failure of ECC is due to all-or-none failure of excitation, other studies suggest failure of excitation is graded. Intracellular recordings of action potentials (APs) in individual fibers during depolarization revealed that APs do not fail in an all-or-none manner. Simultaneous imaging of Ca^2+^ transients during depolarization revealed failure over a narrow range of resting potentials. An AP property that closely correlated with the sudden failure of the Ca^2+^ transient was the integral of AP voltage with respect to time. We hypothesize the close correlation is due to the combined dependence on time and voltage of Ca^2+^ release from the sarcoplasmic reticulum. The quantitative relationships established between resting potential, APs and Ca^2+^ transients provide the foundation for future studies of depolarization-induced failure of ECC in diseases such as periodic paralysis.

## Introduction

The process by which electrical excitation of muscle is converted into force generation is known as excitation contraction coupling (ECC). Successful ECC involves invasion of action potentials into a network of membrane invaginations in muscle known as the transverse tubules (t-tubules) (Adrian et al., 1969). Depolarization in the t-tubules activates Cav1.1 channels, which triggers opening of ryanodine receptors, Ca^2+^ exit from the sarcoplasmic reticulum and force production (Melzer et al., 1995; Dulhunty, 2006; Bannister and Beam, 2013; Hernandez-Ochoa and Schneider, 2018).

Depolarization of the resting membrane potential of skeletal muscle, when severe enough, causes failure of ECC in diseases such periodic paralysis and ICU acquired weakness (Lehmann-Horn et al., 2008; Cannon, 2015; Friedrich et al., 2015) as well as potentially contributing to fatigue during intense exercise (Allen et al., 2008). Studies of depolarization-induced failure of ECC in frog and mammalian skeletal muscle are consistent with all-or-none failure of force generation in individual fibers (Renaud and Light, 1992; Cairns et al., 1997; Cairns et al., 2011). Whole muscle force is generally stable or slightly increased with mild depolarization of the resting potential, which is followed by a steep decline with further depolarization by only a few mV (Holmberg and Waldeck, 1980; Renaud and Light, 1992; Cairns et al., 1997; Yensen et al., 2002; Cairns et al., 2011; Pedersen et al., 2019). The sudden decrease in force is paralleled by a decrease in the Ca^2+^ transient with depolarization of the resting potential beyond -60 mV (Quinonez et al., 2010).

While the idea of all-or-none muscle contraction dates back more than 100 years (Pratt, 1917), the mechanism underlying the near all-or none failure of force generation in the setting of depolarization of the resting potential remains unknown. One hypothesis is that the sudden failure of ECC is due to all-or-none failure of excitability (Renaud and Light, 1992; Cairns et al., 1997). Support for failure of excitability as the mechanism comes from studies showing that the decline in force is paralleled by reduction in extracellular recordings of compound muscle action potentials (Overgaard et al., 1999; Overgaard and Nielsen, 2001; Pedersen et al., 2003). However, intracellular recordings demonstrate graded failure of excitation with depolarization of the resting potential, which manifests as either a gradual reduction in action potential (AP) peak or as APs with variable amplitude that increase with increased current injection (Rich and Pinter, 2001, 2003; Quinonez et al., 2010; Ammar et al., 2015; Miranda et al., 2020; Uwera et al., 2020). Two studies have suggested reduction in AP peak can cause reduction in force generation (Cairns et al., 2003; Gong et al., 2003). These studies are consistent with graded failure of force production in individual fibers.

To determine the mechanism underlying the sudden failure of ECC, we measured muscle force, APs and Ca^2+^ transients in muscle in which the resting potential was depolarized by elevation of extracellular K^+^. In agreement with previous studies, mouse EDL twitch force decreased dramatically over a narrow range of extracellular K^+^ concentrations (Cairns et al., 1997; Yensen et al., 2002). Intracellular recording of APs, combined with Ca^2+^ imaging in individual fibers during elevation of extracellular K^+^, revealed graded failure of excitation was accompanied by relatively sudden failure of the intracellular Ca^2+^ transient. We identified an AP parameter that closely correlated with failure of the Ca^2+^ transient: the area of the AP above -30 mV. Identification of this unexpectedly accurate predictor of failure of the Ca^2+^ transient provides a basis for future studies of depolarization-induced failure of ECC.

## Methods

### Mice

All animal procedures were performed in accordance with the policies of the Animal Care and Use Committee of Wright State University and were conducted in accordance with the United States Public Health Service’s Policy on Humane Care and Use of Laboratory Animals. Mice expressing GCAMP6f (Chen et al., 2013) in skeletal muscle were generated by crossing floxed GCAMP6f mice (Jackson Labs, B6J.Cg-*Gt(ROSA)26Sortm95*.*1(CAG-GCaMP6f)Hze*/MwarJ, cat #028865) with mice expressing parvalbumin promoter driven Cre (Jackson Labs, B6.129P2-Pvalbtm1(cre)Arbr/J, cat# 030218).

### Ex vivo force measurements

Mice were euthanized by inhalation of a saturating dose of isoflurane (∼ 2 g/L) followed by cervical dislocation. The hind limb *extensor digitorum longus* (EDL) muscle was dissected and the proximal tendon of the EDL was tied with a 6-0 caliber silk suture to a bar attached to a custom recording chamber. The distal tendon was tied to a hook and attached to the force transducer (Aurora Scientific). Force measurements were recorded at 21–23°C with the EDL submerged in a solution containing (in mM) 118 NaCl, 3.5 KCl, 1.5 CaCl_2_, 0.7 MgSO_4_, 26.2 NaHCO_3_, 1.7 NaH2PO_4_, and 5.5 glucose and maintained at pH 7.3-7.4 by aeration with 95% O_2_ and 5% CO_2_. Solutions containing elevated concentrations of KCl (3.5, 10, 12, 14, and 16 mM) and with corresponding reduction in NaCl (118, 111.5, 109.5, and 105.5 mM respectively) to maintain a constant osmolarity were used to induce depolarization. The EDL was stimulated with two electrodes placed perpendicularly to the muscle in the bath. The force transducer was controlled by a 305C two-channel controller (Aurora Scientific) and digitized by a Digidata 1550B digitizer (Molecular Devices). A S-900 pulse generator (Dagan) was used to generate 0.1 ms 5V twitch stimulations to the muscle. The pulse generator was triggered using pCLAMP 11 data acquisition and analysis software. The optimal length was determined by adjusting the tension of the muscle until maximal twitch force was achieved. During force recordings, the muscle was exposed to normal K^+^ solution for 20 minutes, followed by high K^+^ solution (10 mM to 16mM) for 45 minutes, and then washed again with normal K^+^ solution for 25 minutes to follow recovery. The EDL was stimulated with a twitch pulse every 5 minutes, and force was recorded.

### Ex vivo recordings of action potentials and Ca^2+^ transients

Mice were sacrificed using CO_2_ inhalation followed by cervical dislocation, and both extensor digitorum longus (EDL) muscles were dissected out tendon-to-tendon. Muscles were maintained and recorded at 22°C within 6 hours of sacrifice. Standard solution contained (in mM): 118 NaCl, 3.5 KCl, 1.5 CaCl_2_, 0.7 MgSO_4_, 26.2 NaHCO_3_, 1.7 NaH2PO_4_, 5.5 glucose, and maintained at pH 7.3-7.4 by aeration with 95% O_2_ and 5% CO_2_.

To prevent contraction, muscles were loaded with 50μM BTS (N-benzyl-p-toluenesulfonamide, Tokyo Chemical Industry, Tokyo, Japan, catalogue #B3082) dissolved in DMSO for 45 minutes prior to recording.

Muscle fibers were impaled with 2 sharp microelectrodes filled with 2 M potassium acetate solution containing 1 mM sulforhodamine 101 (Sigma-Aldrich, Catalogue #S7635) to allow for visualization. Resistances were between 25 and 50 MΩ, and capacitance compensation was optimized prior to recording. APs were evoked by a 0.2 ms injection of current. Fibers with resting potentials more depolarized than –74 mV in solution containing 3.5 mM KCl were discarded.

### Imaging of Ca^2+^ transients

Muscle expressing GCAMP6f was imaged without staining (LeiCa^2+^ I3 cube, band pass 450-490, long pass 515). Imaging was synchronized with triggering of APs using a Master-8 pulse generator (A.M.P.I.). Frames were acquired at 30 frames per second with a sCMOS camera (CS2100M-USB) using ThorCam software (Thorlab Inc. NJ). During infusion of solution containing high K, APs were triggered every 5s. Each AP was synchronized with capture of 48 frames at 30 frame/s. Images were analyzed using Image J (NIH). Regions of interest (ROI) were set both on the muscle fiber being stimulated and a neighboring fiber. The neighboring fiber was used to record background, which was subtracted from the ROI on the stimulated fiber.

### Fitting of data with Boltzmann distributions

Data for AP peak vs. resting potential, Ca^2+^ image intensity vs. AP, and Ca^2+^ image intensity vs. resting potential were all fit to a Boltzmann, 

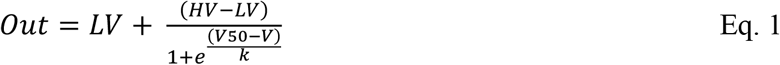

where *Out* represents the dependent variable (either AP peak or Ca^2+^ image intensity), *V* is the independent voltage variable (either resting potential or AP peak), *LV* is the limiting value when *V* is very low (toward more negative), *HV* is the limiting value when *V* is very high (toward more positive), *V50* is the value of *V* at which *Out* is halfway between *HV* and *LV*, and k is the slope factor. All voltages and the variable k are expressed in mV, and Ca^2+^ image intensity is in arbitrary units between 1 for maximum intensity for each experiment and 0.

### Statistics

Data for recordings from different muscles were analyzed using nested analysis of variance with n as the number of mice, with data presented as mean ± SD. Comparisons of different parameters recorded from the same fiber were compared using the paired students t-test. *p* < 0.05 was considered to be significant. The numbers of animals and fibers for comparisons are described in the corresponding figure legends and text.

## Results

Measurement of twitch force in the mouse EDL following elevation of extracellular K^+^ to between 10 and 16 mM yielded results similar to those previously reported. There was an initial increase in force (Fig 1A, n= 3 muscles for each K^+^ concentration) (Cairns et al., 1997; Yensen et al., 2002; Pedersen et al., 2019). The initial increase in force is paralleled by an increase in the Ca^2+^ transient, which occurs despite a reduction of AP peak (Quinonez et al., 2010; Pedersen et al., 2019). The mechanism underlying the increase in Ca^2+^ transient remains unknown; it does not appear to be due to changes in characteristics of APs (Yensen et al., 2002).

**Figure 1:**
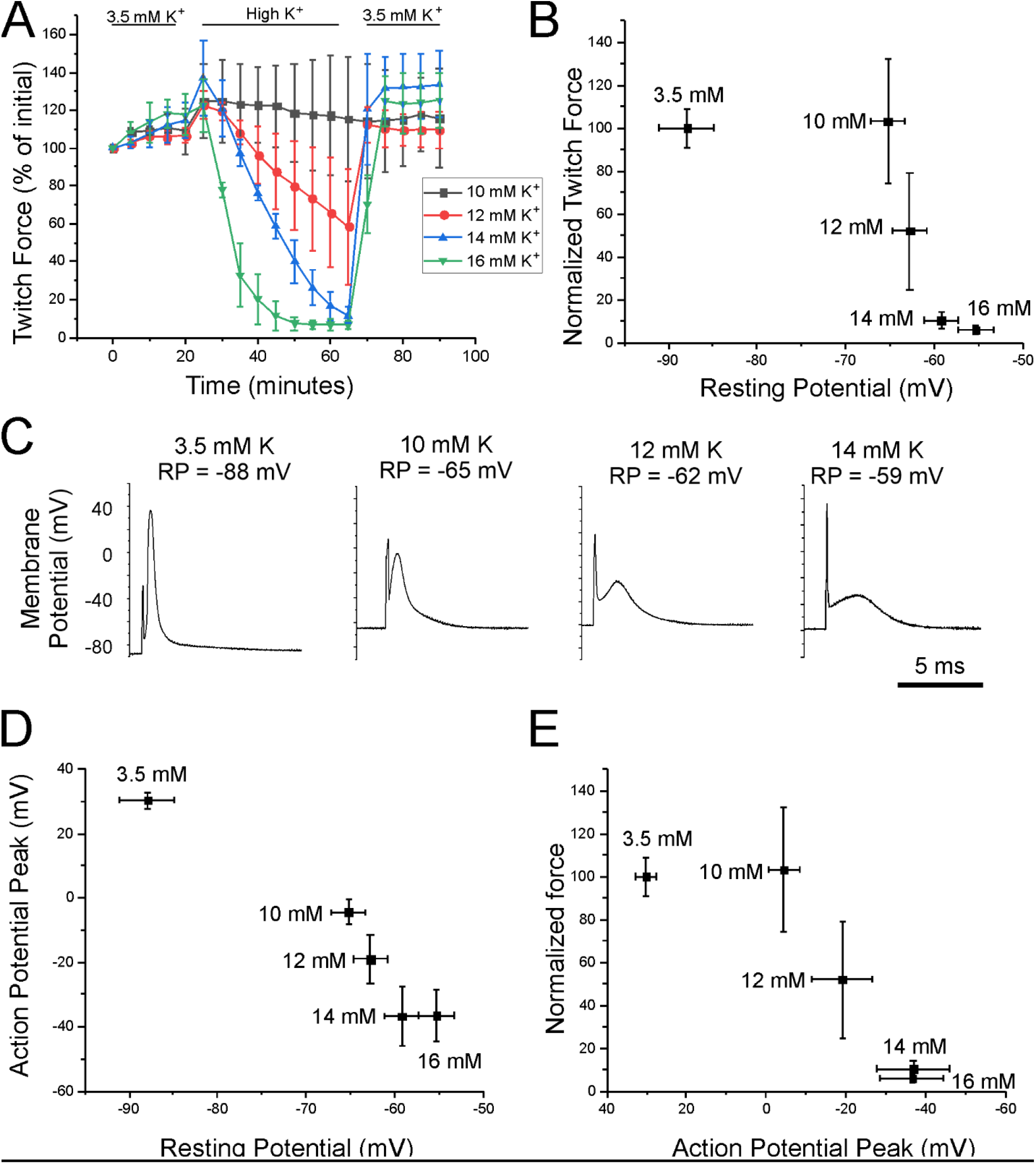
Loss of force secondary to depolarization of the resting potential is accompanied by graded failure of excitation. A) Shown is a plot of extensor digitorum longus (EDL) twitch force versus time following infusion of various concentrations of external K^+^ (n = 3 muscles for each K^+^ concentration, error bars = SD). Force for each muscle was normalized to the initial force recorded. B) Mean force after 40 minutes in high K^+^ (from Fig 1A) normalized to force after 40 minutes in 3.5 mM K^+^ plotted versus mean resting membrane potential recorded 20 to 40 minutes following infusion of high K^+^ solution. C) Examples of typical APs recorded at various resting membrane potentials. In each trace the AP is preceded by a 0.2 ms stimulus artifact. D) Plot of mean AP peak versus mean resting potential for each of the K^+^ concentrations. E) Plot of mean force 40 minutes following elevation of K^+^ versus mean AP peak for each K^+^ concentration.

Following the initial increase in force, there was a decline that became faster with higher levels of extracellular K^+^ (Fig 1A). With return to solution containing normal K^+^, force recovered rapidly. Force generated 40 minutes following infusion of high K^+^ was steeply dependent on extracellular K^+^, such that force was near normal in 10 mM K^+^ but near 0 in 14 and 16 mM K^+^. The mean resting potential 20-40 minutes after infusion of each concentration of K^+^was measured in a separate set of experiments (n = 80 fibers from 4 muscles for each K^+^ concentration) and those data were used to construct a plot of force versus mean resting potential. There was a steep loss of force over a narrow range of resting potentials: normal force was generated at a mean resting potential of -65.3 ± 1.9 mV in 10 mM K^+^ and almost no force was generated at a mean resting potential of -59.3 ± 1.9 mV in 14 mM K^+^ (Fig 1B). Our finding is similar to a previous report of steep dependence of force production on resting potential (Cairns et al., 1997).

The sudden loss of force with depolarization of muscle has been hypothesized to be due to all-or-none failure of APs (Renaud and Light, 1992; Cairns et al., 1997). To test this hypothesis, we measured intracellular APs from EDL fibers in each of the high K^+^ solutions. APs were measured with a voltage-sensing electrode and triggered with a current-passing electrode. While APs became smaller with depolarization, responses were still present in 14 and16 mM K^+^; concentrations of K^+^ in which force production is near 0 (Fig 1B, C). These data counter the idea of all-or-none APs, and agree with several studies suggesting more gradual failure of excitation (Rich and Pinter, 2001, 2003; Quinonez et al., 2010; Ammar et al., 2015; Miranda et al., 2020; Uwera et al., 2020).

If depolarization of the resting potential triggers a gradual decline of APs, why is there close to all-or-none failure of force production with depolarization? One possibility is that small APs fail to trigger elevation of intracellular Ca^2+^. At resting potentials between -80 and -90 mV, APs peaked at 30.2 ± 2.7 mV. With depolarization of the resting potential to -65.3 ± 1.9 mV, the peak was reduced to -4.4 ± 3.9 mV and at a resting potential of -55.4 ± 2.0 mV, the peak averaged -36.5 ± 8.0 mV (Fig 1D). Reduction of the mean AP peak from 30.2 mV to -4.4 mV was associated with little to no reduction in force, whereas reduction of the peak from -4 mV to - 36 mV was associated with almost complete loss of force (Fig 1 E). These data raise the possibility that there may be a threshold for AP peak above which APs trigger full contraction and below which there is failure of ECC.

The relationship between depolarization of the resting potential and reduction in AP peak has never been studied in individual fibers. To determine the contribution of damage-induced depolarization to AP changes during prolonged impalement, we performed 7-minute recordings of resting potential and AP peaks in solution containing normal external K^+^. Seven minutes after impalement, impalement-induced damage caused 8 to 20 mV of depolarization of the resting membrane potential (Fig 2 A, B), which was accompanied by up to 15 mV of reduction in AP peak (Fig 2 B). Infusion of a solution containing 16 mM K^+^ causes substantial depolarization of the resting membrane potential and reduction in the AP peak beyond that seen with impalement alone (Fig 2 C, D). Consistent with recordings taken from populations of fibers, infusion of 16 mM K^+^ caused graded reduction of the AP peak from a maximum ranging from +15 to +35 mV down to -25 to -45 mV. Between resting potentials of -65 to -52 mV, there was rapid reduction of the peak with further depolarization. To quantitate the relationship between depolarization of the resting potential and reduction in AP peak, we fit the data for individual fibers (plots in Fig 2D) with Boltzmann equations (Eq. 1, See Fig 3G for an example fit). For these fits, the HV limit, which represents the minimal AP peak when resting potential was elevated, was constrained to be between -30 mV and -50 mV. The V50 for the resting potential at which AP peak was half maximal was -58.2 ± 3.3 mV, the slope factor *k* was 1.8 ± 0.6 mV, and the average value of the half-maximal AP peak at the V50 value was -15.5 ± 4.9 mV (n =12 fibers from 6 mice).

**Fig 2:**
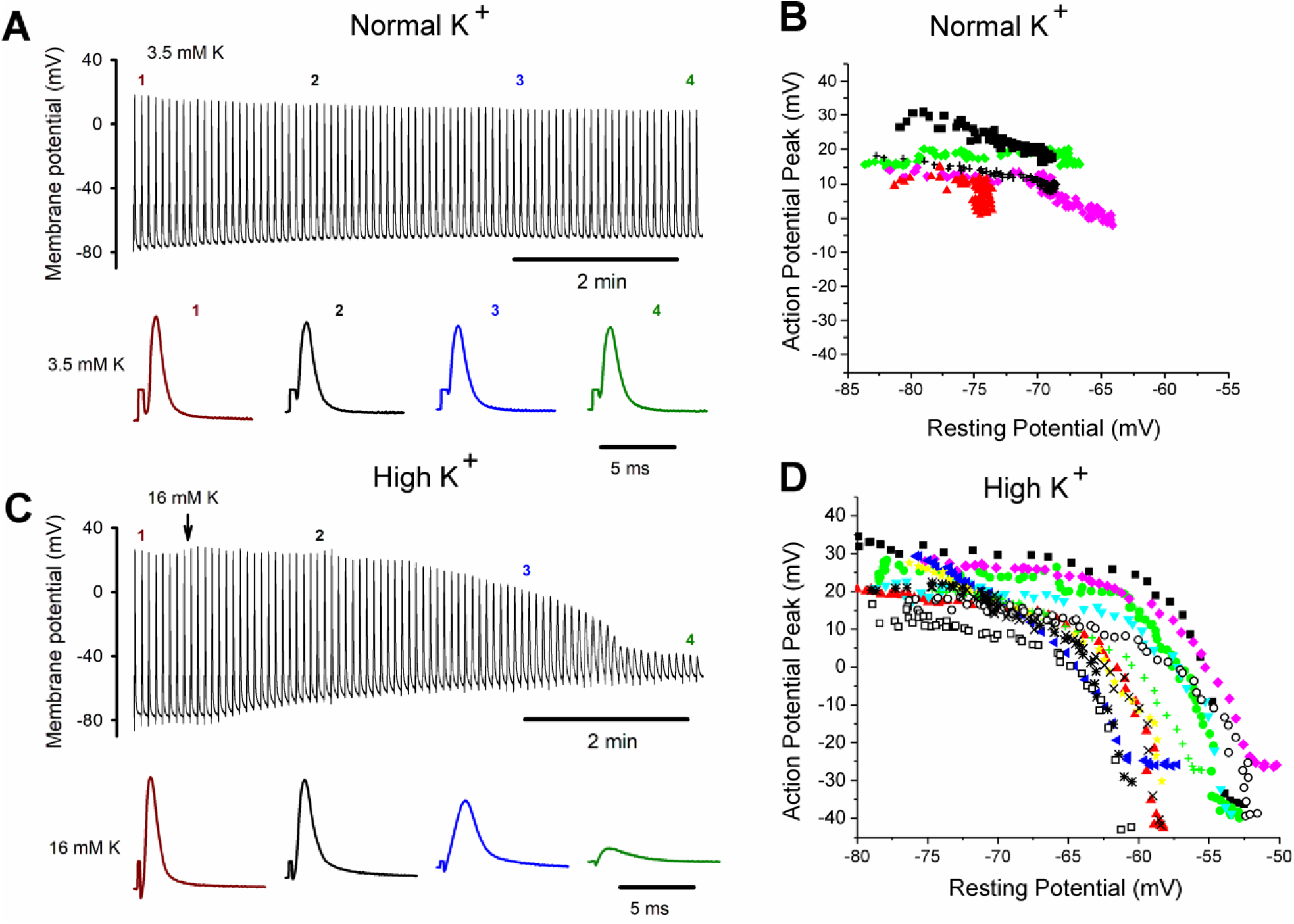
Failure of excitation in individual fibers during depolarization. A) Top: Shown are APs from a fiber during a 7-minute recording in 3.5 mM K^+^. APs were triggered at 0.2 Hz by a 0.2 ms injection of depolarizing current of constant amplitude that was 150% of initial threshold current. The recording is not continuous: a 5 ms block of time is shown for each AP and stimulus artifacts have been removed. The time base indicated is for the time between APs. Bottom: Shown on an expanded time base are APs from the top traces at the time points indicated by the numbers. Stimulus artifacts have been truncated for clarity. B) Plot of AP peak versus resting potential during for 5 fibers in which K^+^ was maintained at 3.5 mM throughout the 7-minute recording. The impalement-induced depolarization in the 5 fibers ranged from 8 to close to 20 mV. C) Top: An intracellular recording from a fiber during infusion of 16 mM K^+^. The infusion began at the time indicated by the vertical arrow. Bottom: Shown on an expanded time base are APs from the time points indicated by the numbers. D) Plot of AP peak versus resting potential during infusion of 16 mM K^+^ for 12 fibers.

**Figure 3:**
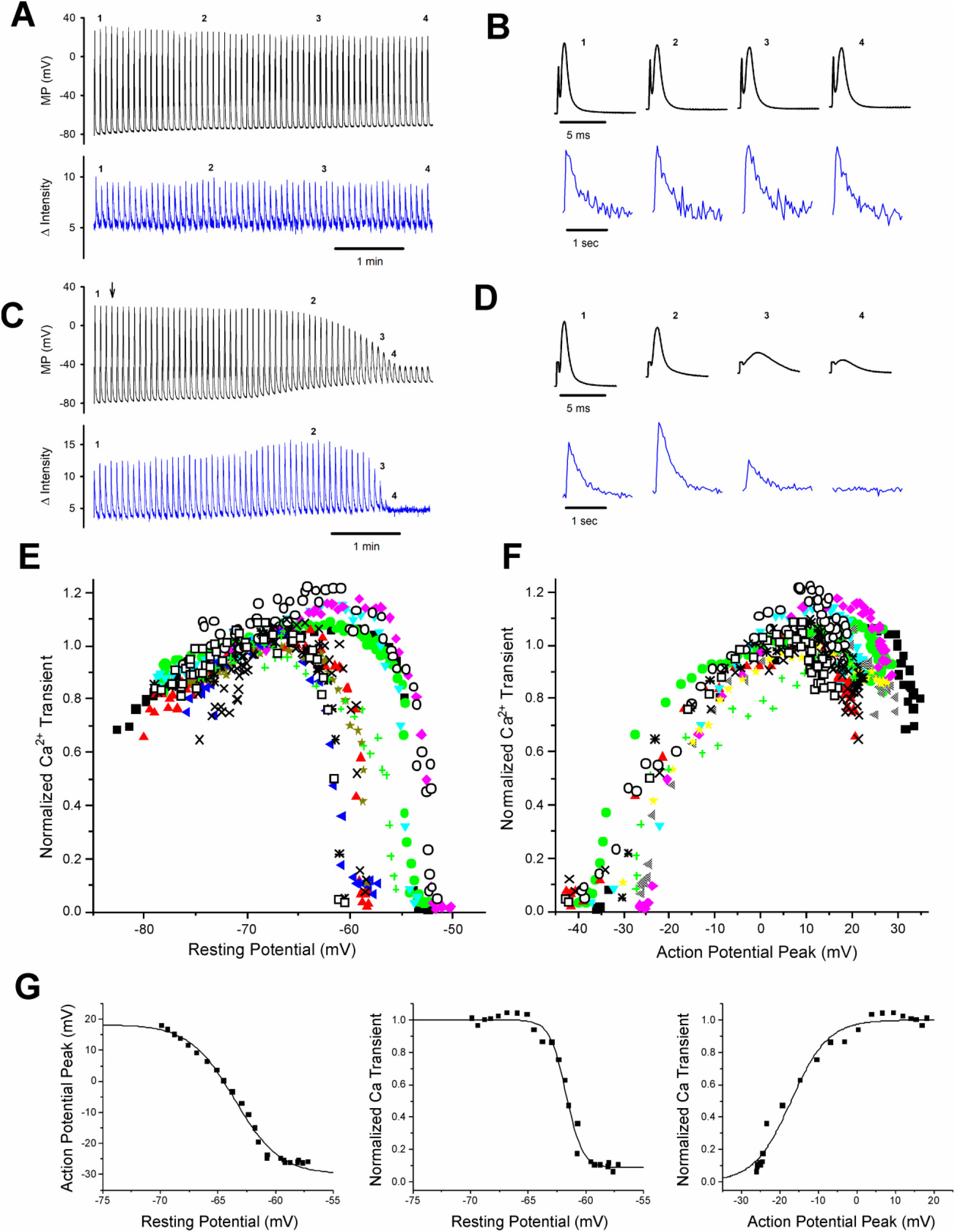
Failure of the Ca^2+^ transient with depolarization. A) Shown are the AP and Ca^2+^ transient for a fiber expressing GCAMP6f in 3.5 mM K^+^ during a recording. The stimulus artifact in the AP traces has been eliminated for clarity. The recording is not continuous. A 5 ms block of time is shown for each AP and a 1 s block of time is shown for each Ca^2+^ transient. The time base indicated is for the time between APs and Ca^2+^ transients. B) Shown on an expanded time scale are the APs and corresponding Ca^2+^ transients for the 4 time points indicated in A. C) APs and Ca^2+^ transients for a fiber during infusion of solution containing 16 mM K^+^. D) Shown on an expanded time scale are APs and corresponding Ca^2+^ transients for the 4 time points indicated in C. E) Plot of the peak of the Ca^2+^ transient versus resting potential for the 12 fibers studied (The same fibers studied in Fig 2). The peak of the Ca^2+^ transient present at a resting potential of -70 mV was normalized to a value of 1 for each fiber. F) Plot of normalized Ca^2+^ transient versus AP peak for the 12 fibers studied. G) Shown are example Boltzmann fits of AP Peak and Ca^2+^ transient versus resting potential as well as Ca^2+^ transient versus AP peak for a single fiber.

In order to better understand how depolarization of the resting potential affects generation of force, we recorded APs while simultaneously imaging Ca^2+^ transients in fibers from mice expressing GCAMP6f in skeletal muscle. GCAMP6f is a high affinity genetically-encoded Ca^2+^ indicator which allowed us to determine changes in amplitude of the Ca^2+^ transient even during low release flux events, but which does not allow for examination of kinetics (Chen et al., 2013; Shang et al., 2014; Dana et al., 2019). In normal extracellular K^+^, single APs generated a bright fluorescent transient (Fig 3A-D, video associated with this submission). When extracellular K^+^ was kept at 3.5 mM, the peak of the Ca^2+^ transient was stable over time, with a slight trend towards increasing (Fig 3A, B, 1.16 at 7 minutes vs initial value normalized to 1, p =.07, n = 5).

With depolarization during infusion of 16 mM K^+^, there was an initial increase in the Ca^2+^ transient from a mean normalized value of 0.81 ± 0.10 to the normalized maximum of 1 (p < .0001 paired t-test, n =12, Fig 3C-E). This increase is similar to what has been reported previously (Quinonez et al., 2010; Pedersen et al., 2019) and occurred despite reduction in AP peak (Fig 3D, see Fig 2D for AP peak plots for the same 12 fibers). As depolarization progressed there was rapid, complete loss of the Ca^2+^ transient (Fig 3C-E). To quantitate the relationship between depolarization and failure of the Ca^2+^ transient we fit the data for the decrease in Ca^2+^ transient with depolarization of the resting potential beyond -70 mV with a Boltzmann equation (Eq. 1, See Fig 3G for an example fit). For these fits, the LV limit, which represents the Ca^2+^ image intensity when resting potential was -70 mV, was fixed to 1, and the HV limit was constrained to be between 0 and 0.1. The resting potential at which Ca^2+^ transient was half maximal was -57.5 ± 3.4 mV with a slope factor of 0.4 ± 0.2 mV, which was significantly steeper than the slope for reduction in AP peak (p < 1 × 10^−5^, paired t-test, n =12 fibers from 6 mice, Fig 3E).

We plotted the reduction in Ca^2+^ transient versus AP peak (Fig 3F), and fit the data with a Boltzmann equation (Eq. 1, see Fig 3G for an example fit). For these fits, the HV was fixed to 1 and the LV was fixed to 0. The Ca^2+^ transient was half maximal at an AP peak of -21.0 ± 4.0 mV with a slope factor of 5.9 ± 1.7 mV. This relationship between peak voltage and Ca^2+^ transient was within the range of values obtained from voltage clamp studies of mouse muscle fibers (Wang et al., 1999; Gregorio et al., 2017). These data suggest APs peaking below -30 mV trigger little to no elevation of intracellular Ca^2+^ and hence little to no generation of force.

The finding that all-or none-failure of the AP was not the mechanism underlying depolarization-induced failure of ECC raises the possibility that measurement of APs might not allow for accurate prediction of successful ECC. This would necessitate Ca^2+^ imaging to determine the efficacy of APs in triggering ECC. We wished to identify an AP property that would accurately predict the Ca^2+^ transient. We first examined the correlation between normalized AP amplitude and the Ca^2+^ transient. A loss of 11.6 ± 1.7 mV of resting potential was required to reduce AP amplitude from 90% to 10% of maximum (Fig 4D). In contrast, the Ca^2+^ transient was reduced from 90% to 10% of maximum with a loss of only 4.1 ± 2.4 mV of resting potential (Fig 4D, p < 1×10^−6^ vs APs, n=12, paired student’s t-test). This statistically significant difference led us to look for another AP parameter that decreased more sharply with depolarization of the resting potential.

**Fig 4.**
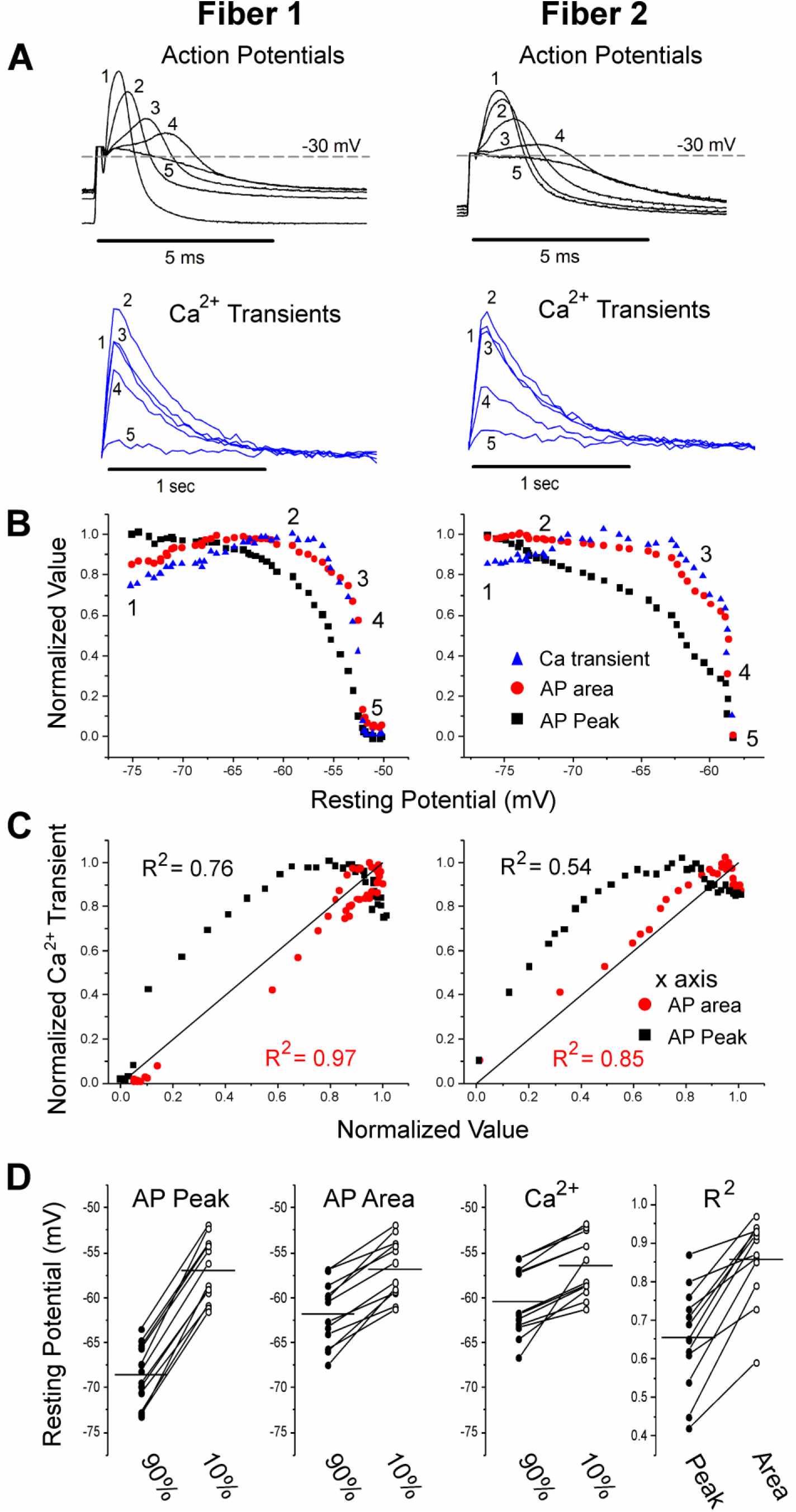
Correlation between Ca^2+^ transient and AP area. A) Shown in black are the AP traces and Ca^2+^ transients recorded from 2 muscle fibers during infusion of 16 mM K+. A horizontal line at -30 mV represents the cutoff for measurement of AP area and normalized AP peak. APs peaking below -30 mV had peaks and areas set to 0. Below the AP traces shown in blue are the corresponding Ca^2+^ transients. The stimulus artifact has been truncated in the AP traces for clarity. B) Plots of normalized AP peak, normalized AP area and normalized Ca^2+^ transient versus resting potential for 2 fibers. The numbers 1-5 on each plot represent the points corresponding to the 5 AP and Ca^2+^ transient traces shown in A. C) Plots of the normalized Ca^2+^ transient versus either AP area or AP peak for the 2 fibers. The line of identity is drawn on each plot as a reference. The R^2^ value for each relationship is shown on the graph. D) Plots of the resting potential at which AP peak, AP area and the Ca^2+^ transients are 90% and 10% of maximal as well as the plot of the R^2^ values for the relationship between AP peak and AP area vs the Ca^2+^ transient for each of the 12 fibers. The horizontal line in each plot represents the mean value for the data.

As shown in Fig 3F, an AP peak above -30 mV is required to consistently trigger Ca^2+^ release. We thus set AP peaks of -30 mV or below to 0 and normalized AP amplitude. At mildly depolarized resting potentials, drops in normalized AP peaks were accompanied by increases in the Ca^2+^ transient (Fig 4A, B, Fig 3E) (Quinonez et al., 2010; Pedersen et al., 2019). At more depolarized resting potentials, the drop in AP peak did not closely track the sharp drop in Ca^2+^ transient (Fig 4A, B). When the normalized Ca^2+^ transient was plotted against the normalized AP peak, the mean R^2^ was 0.65 ± 0.14 (Fig 4C, D, n =12 fibers). This was not the close relationship we were hoping to find.

We next considered whether changes in AP kinetics might affect the Ca^2+^ transient. It has been shown previously that AP half width increases with depolarization of the resting membrane potential due to both decreased rate of rise and decreased rate of decay of the AP (Yensen et al., 2002). It has also been shown previously that increasing AP half width by blocking K^+^ channels increases twitch force (Delbono and Kotsias, 1987; van Lunteren et al., 2006), presumably secondary to increases in the Ca^2+^ transient. It is thus possible that increases in AP half width lessen the effect of decreasing AP peak on the Ca^2+^ transient during modest depolarization of the resting potential.

To include changes in both AP half width and peak, we measured the integral of AP voltage with respect to time. The integral above -30 mV was used to account for the lack of Ca^2+^ transient when APs peaked below -30 mV. AP area closely paralleled Ca^2+^ transient during depolarization of the resting potential (Fig 4A, B). When the normalized Ca^2+^ transient was plotted against normalized AP area, the mean R^2^ value was 0.86 ± 0.11 (Fig 4C, D, p < .001 vs the R^2^ value for Ca^2+^ transient vs AP peak, n = 12 fibers, paired t-test). The R^2^ value was larger for AP area vs Ca^2+^ transient because AP area closely mimicked the rapid decrease from 90% to 10% of maximum that occurred for the Ca^2+^ transient (Fig 4D, p = 0.27 vs the range of resting potentials for the decrease in Ca^2+^ transient, paired students t-test). These data suggest measurement of AP area is a significantly better predictor of failure of the Ca^2+^ transient than AP peak.

## Discussion

Prolonged intracellular recordings of individual muscle fibers during infusion of solution containing elevated K^+^ were performed while simultaneously imaging the Ca^2+^ transient triggered by action potentials (APs). The sudden failure of ECC with depolarization of the resting potential was not due to all-or-none failure of APs. An AP property that closely correlated with Ca^2+^ transient and thus the sudden failure of ECC was the integral of AP voltage with respect to time. Ours is the first study to determine quantitative relationships between resting potential, APs and the Ca^2+^ transients that trigger contraction during ECC in skeletal muscle. Understanding these relationships provides the foundation for future studies of depolarization-induced failure of ECC in various disease states such as periodic paralysis.

### The integral of AP voltage with respect to time as the determinant of the Ca^2+^ transient

In this and a previous study, it was found that during elevation of extracellular K^+^, reduction in AP peaks occurs over a relatively wide range of resting potentials (Ammar et al., 2015). The decrease is relatively modest between resting potentials of -85 to – 65 mV, but becomes rapid between -65 mV and -55mV. This is due to the non-linear relationship between the density of Na^+^ channels available to participate in APs and AP peak (Rich and Pinter, 2001, 2003). The lack of all-or-none AP failure raised the question of why failure of muscle force generation is so sudden, occurring over a narrow range of resting potentials (Holmberg and Waldeck, 1980; Renaud and Light, 1992; Cairns et al., 1997; Yensen et al., 2002; Cairns et al., 2011; Pedersen et al., 2019). Ca^2+^ imaging during depolarization revealed the answer: there is failure of the Ca^2+^ transient over a narrow range of resting potentials.

Our study and previous studies suggest Ca^2+^ release from the sarcoplasmic reticulum begins to be triggered at voltages of -30 to -20 mV and becomes maximal at voltages above +10 mV (Wang et al., 1999; Braubach et al., 2014). Because APs peaking below -30 mV do not trigger elevation of Ca^2+^, we set -30 mV as 0 and normalized AP peaks. Despite normalization, the relationship between AP peak and Ca^2+^ transient was not as close as we had hoped: the reduction in the AP peak still occurred more gradually than the drop in Ca^2+^ transient. This caused us to look for a measure of APs that would more closely correlate with the Ca^2+^ transient.

It has previously been reported that depolarization of the resting potential triggers widening of APs (Yensen et al., 2002). To incorporate consideration of both peak voltage and AP half width into our measure of APs, we took the integral of AP voltage with respect to time. This measure of APs closely correlated with the decrease in Ca^2+^ transient responsible for failure of ECC secondary to depolarization of the resting potential. AP area did not correlate with the increase in Ca^2+^ transient occurring with mild depolarization of the resting potential. There is still a factor yet to be determined that is responsible for the initial increase in Ca^2+^ transient with depolarization of the resting potential.

The finding that AP integral more closely correlates with failure of the Ca^2+^ transient than did AP peak makes biophysical sense. Depolarization during an AP triggers elevation of intracellular Ca^2+^ by causing movement of gating charges in Cav1.1 channels in the t-tubules (Kovacs et al., 1979; Rios and Brum, 1987; Garcia et al., 1994). The movement of the voltage-sensing particles cause further conformational rearrangements within Ca_V_1.1 which open RyR1 and provide a conduit for release of Ca^2+^ from the sarcoplasmic reticulum into the myoplasm. The total gating charge is dependent on both voltage and time (Schneider and Chandler, 1973). Thus, the longer the membrane potential remains depolarized during an AP, the greater the gating charge movement produced by Cav1.1 channels until saturation is reached. Established data on Cav1.1 gating charge movement indicates that both the magnitude and time-course of gating charge movement in response to membrane potential changes are complex and non-linear (Gregorio et al., 2017). The release of Ca^2+^ in response to Cav1.1 gating charge movement is also non-linear. The use of the AP area metric as described in this work is therefore a simplification of the underlying processes.

There are several limitations of our study. One is that the Ca^2+^ indicator used is relatively high affinity (Chen et al., 2013; Shang et al., 2014; Dana et al., 2019), such that saturation of the dye may have caused us to underestimate the magnitude of the Ca^2+^ transient. One factor that might contribute to the sudden failure of the Ca^2+^ transient independent of AP area, is inactivation of Ca^2+^ release (Cota et al., 1984; Chua and Dulhunty, 1988; Schneider and Simon, 1988; Robin and Allard, 2013; Gregorio et al., 2017; Hernandez-Ochoa and Schneider, 2018). The voltage dependence of Cav1.1 gating charge availability has been measured and found to have a midpoint of -57 mV (Gregorio et al., 2017); the same membrane potential at which we found the Ca^2+^ transient was half maximal. We did not separate the effect of depolarization of the resting potential on APs from the effect of depolarization of resting potential on Ca^2+^ release from the sarcoplasmic reticulum. Finally, we chose a voltage cut off for the integral of AP of -30 mV. This may not be the correct cut off for all fibers. As shown in Fig 3F. there was a 15 mV range of AP peaks in different fibers at which there began to be a Ca^2+^ transient. Despite all the caveats, the close correlation between AP area and Ca^2+^ transient suggests the approach has value and may prove useful in future studies of ECC.

### The definition of an AP

Textbooks often describe APs as sudden depolarizations, which are all-or-none (Boron and Boulpaep, 2017; Purves et al., 2018). This definition derives from the finding that AP amplitude and conduction are independent of the amount of current injected (Cole and Curtis, 1939). APs triggered at a normal muscle resting potential of -80 to -85 mV are all-or-none. However, with depolarization of the resting potential there is graded reduction of the AP peak such that AP amplitude ranges from 120 mV to below 10 mV (the current study and (Rich and Pinter, 2003; Quinonez et al., 2010; Ammar et al., 2015; Miranda et al., 2020; Uwera et al., 2020)).

If APs are not all-or-none, how does one define what is and what isn’t an AP? One definition might be transient depolarizations that propagate the length of the fiber. However, to use this definition propagation of APs along the length of the fiber must be measured, which although possible, is not trivial (Riisager et al., 2014). As we did not study AP propagation, our study does not shed light on whether failure of AP propagation contributes to failure of ECC with depolarization of the resting potential.

A second definition of APs could be depolarizations that trigger elevation of intracellular Ca^2+^. The focus on elevation of Ca^2+^ is appealing since it is the link between APs and ECC. With this definition, the presence of an AP would always correlate with successful ECC. However, this definition requires concurrent imaging of intracellular Ca^2+^, which limits applicability.

A recent study defined inexcitable fibers as fibers, “for which the membrane potential did not change at all following a stimulation” (Uwera et al., 2020). Although not explicitly stated, a corollary of this definition is that any depolarization occurring after termination of a stimulation, no matter how small, is an AP. With this definition, APs can vary widely in amplitude, ability to conduct along the length of the fiber, and ability to trigger ECC. The lack of functional correlation is a weakness, but this definition avoids making binary decisions about what is and what isn’t an AP, which our study suggests would be difficult. Thus, despite its limitations, we favor this definition for studies of muscle diseases with failure of ECC caused by depolarization of the resting potential.

### Slow Inactivation of Na channels likely contributes to depolarization-induced failure of ECC

Recording in individual fibers during infusion of 16 mM K^+^ resulted in higher AP peaks at a given resting potential than were obtained from sampling fibers from muscles perfused with different concentrations of K^+^ (Fig 1D vs Fig 2D). One potential explanation of this difference is the speed of depolarization. In recordings from individual fibers, infusion of 16 mM K^+^ triggered a ∼30 mV depolarization over several minutes. When sampling fibers from muscles in solutions with varying K^+^ concentrations, muscles were incubated in each solution for 20 minutes prior to recording of APs. The prolonged depolarization allows for greater slow inactivation of Na^+^ channels, (Ruff, 1999; Rich and Pinter, 2003), such that AP peak decreased at more negative resting potentials. In disease states causing depolarization of the resting potential, depolarization is generally prolonged such that slow inactivation of Na^+^ channels would play an important role in the reduction of AP peak.

### Summary

Ours is the first study to establish quantitative relationships between resting potential, APs and generation of the Ca^2+^ transient in individual fibers. Understanding these relationships provides the foundation for studies of depolarization-induced failure of ECC in a variety of muscle diseases.

## References

Adrian RH, Costantin LL, Peachey LD (1969) Radial spread of contraction in frog muscle fibres. J Physiol 204:231–257.

Allen DG, Lamb GD, Westerblad H (2008) Skeletal muscle fatigue: cellular mechanisms. Physiol Rev 88:287–332.

Ammar T, Lin W, Higgins A, Hayward LJ, Renaud JM (2015) Understanding the physiology of the asymptomatic diaphragm of the M1592V hyperkalemic periodic paralysis mouse. J Gen Physiol 146:509–525.

Bannister RA, Beam KG (2013) Ca(V)1.1: The atypical prototypical voltage-gated Ca(2)(+) channel. Biochim Biophys Acta 1828:1587–1597.

Boron WF, Boulpaep EL (2017) Medical Physiology, Third Edition. Philadelphia, PA: Elsevier. Braubach P, Orynbayev M, Andronache Z, Hering T, Landwehrmeyer GB, Lindenberg KS, Melzer W (2014) Altered Ca(2+) signaling in skeletal muscle fibers of the R6/2 mouse, a model of Huntington’s disease. J Gen Physiol 144:393–413.

Cairns SP, Leader JP, Loiselle DS (2011) Exacerbated potassium-induced paralysis of mouse soleus muscle at 37 degrees C vis-a-vis 25 degrees C: implications for fatigue. K+ - induced paralysis at 37 degrees C. Pflugers Arch 461:469–479.

Cairns SP, Buller SJ, Loiselle DS, Renaud JM (2003) Changes of action potentials and force at lowered [Na+]o in mouse skeletal muscle: implications for fatigue. Am J Physiol Cell Physiol 285:C1131–1141.

Cairns SP, Hing WA, Slack JR, Mills RG, Loiselle DS (1997) Different effects of raised [K+]o on membrane potential and contraction in mouse fast- and slow-twitch muscle. Am J Physiol 273:C598–611.

Cannon SC (2015) Channelopathies of skeletal muscle excitability. Compr Physiol 5:761–790.

Chen TW, Wardill TJ, Sun Y, Pulver SR, Renninger SL, Baohan A, Schreiter ER, Kerr RA, Orger MB, Jayaraman V, Looger LL, Svoboda K, Kim DS (2013) Ultrasensitive fluorescent proteins for imaging neuronal activity. Nature 499:295–300.

Chua M, Dulhunty AF (1988) Inactivation of excitation-contraction coupling in rat extensor digitorum longus and soleus muscles. J Gen Physiol 91:737–757.

Cole KS, Curtis HJ (1939) Electric Impedance of the Squid Giant Axon during Activity. J Gen Physiol 22:649–670.

Cota G, Nicola Siri L, Stefani E (1984) Calcium channel inactivation in frog (Rana pipiens and Rana moctezuma) skeletal muscle fibres. J Physiol 354:99–108.

Dana H, Sun Y, Mohar B, Hulse BK, Kerlin AM, Hasseman JP, Tsegaye G, Tsang A, Wong A, Patel R, Macklin JJ, Chen Y, Konnerth A, Jayaraman V, Looger LL, Schreiter ER, Svoboda K, Kim DS (2019) High-performance calcium sensors for imaging activity in neuronal populations and microcompartments. Nat Methods 16:649–657.

Delbono O, Kotsias BA (1987) Relation between action potential duration and mechanical activity on rat diaphragm fibers. Effects of 3,4-diaminopyridine and tetraethylammonium. Pflugers Arch 410:394–400.

Dulhunty AF (2006) Excitation-contraction coupling from the 1950s into the new millennium. Clin Exp Pharmacol Physiol 33:763–772.

Friedrich O, Reid MB, Van den Berghe G, Vanhorebeek I, Hermans G, Rich MM, Larsson L (2015) The Sick and the Weak: Neuropathies/Myopathies in the Critically Ill. Physiol Rev 95:1025–1109.

Garcia J, Tanabe T, Beam KG (1994) Relationship of calcium transients to calcium currents and charge movements in myotubes expressing skeletal and cardiac dihydropyridine receptors. J Gen Physiol 103:125–147.

Gong B, Legault D, Miki T, Seino S, Renaud JM (2003) KATP channels depress force by reducing action potential amplitude in mouse EDL and soleus muscle. Am J Physiol Cell Physiol 285:C1464–1474.

Gregorio JF, Pequera G, Manno C, Rios E, Brum G (2017) The voltage sensor of excitation-contraction coupling in mammals: Inactivation and interaction with Ca2+. Journal of General Physiology 149:1041–1058.

Hernandez-Ochoa EO, Schneider MF (2018) Voltage sensing mechanism in skeletal muscle excitation-contraction coupling: coming of age or midlife crisis? Skelet Muscle 8:22.

Holmberg E, Waldeck B (1980) On the possible role of potassium ions in the action of terbutaline on skeletal muscle contractions. Acta Pharmacol Toxicol (Copenh) 46:141–149.

Kovacs L, Rios E, Schneider MF (1979) Calcium transients and intramembrane charge movement in skeletal muscle fibres. Nature 279:391–396.

Lehmann-Horn F, Jurkat-Rott K, Rudel R (2008) Diagnostics and therapy of muscle channelopathies--Guidelines of the Ulm Muscle Centre. Acta Myol 27:98–113.

Melzer W, Herrmann-Frank A, Luttgau HC (1995) The role of Ca2+ ions in excitation-contraction coupling of skeletal muscle fibres. Biochim Biophys Acta 1241:59–116.

Miranda DR, Reed E, Jama A, Bottomley M, Ren H, Rich MM, Voss AA (2020) Mechanisms of altered skeletal muscle action potentials in the R6/2 mouse model of Huntington’s disease. Am J Physiol Cell Physiol.

Overgaard K, Nielsen OB (2001) Activity-induced recovery of excitability in K(+)-depressed rat soleus muscle. Am J Physiol Regul Integr Comp Physiol 280:R48–55.

Overgaard K, Nielsen OB, Flatman JA, Clausen T (1999) Relations between excitability and contractility in rat soleus muscle: role of the Na+-K+ pump and Na+/K+ gradients. J Physiol 518:215–225.

Pedersen KK, Cheng AJ, Westerblad H, Olesen JH, Overgaard K (2019) Moderately elevated extracellular [K(+)] potentiates submaximal force and power in skeletal muscle via increased [Ca(2+)]i during contractions. Am J Physiol Cell Physiol 317:C900–C909.

Pedersen TH, Clausen T, Nielsen OB (2003) Loss of force induced by high extracellular [K+] in rat muscle: effect of temperature, lactic acid and beta2-agonist. J Physiol 551:277–286.

Pratt FH (1917) The all-or-none principle in graded response of muscle contraction. American Journal of Physiology 44:517–542.

Purves D, Augustine GJ, Fitzpatrick D, Hall WC, S. La, Mooney RD, Platt ML, White L (2018) Neuroscience. New York, New York: Oxford University Press.

Quinonez M, Gonzalez F, Morgado-Valle C, DiFranco M (2010) Effects of membrane depolarization and changes in extracellular [K(+)] on the Ca (2+) transients of fast skeletal muscle fibers. Implications for muscle fatigue. J Muscle Res Cell Motil 31:13–33.

Renaud JM, Light P (1992) Effects of K+ on the twitch and tetanic contraction in the sartorius muscle of the frog, Rana pipiens. Implication for fatigue in vivo. Can J Physiol Pharmacol 70:1236–1246.

Rich MM, Pinter MJ (2001) Sodium channel inactivation in an animal model of acute quadriplegic myopathy. Annals of Neurology 50:26–33.

Rich MM, Pinter MJ (2003) Crucial role of sodium channel fast inactivation in muscle fibre inexcitability in a rat model of critical illness myopathy. J Physiol 547:555–566.

Riisager A, Duehmke R, Nielsen OB, Huang CL, Pedersen TH (2014) Determination of cable parameters in skeletal muscle fibres during repetitive firing of action potentials. J Physiol 592:4417–4429.

Rios E, Brum G (1987) Involvement of dihydropyridine receptors in excitation-contraction coupling in skeletal muscle. Nature 325:717–720.

Robin G, Allard B (2013) Major contribution of sarcoplasmic reticulum Ca(2+) depletion during long-lasting activation of skeletal muscle. J Gen Physiol 141:557–565.

Ruff RL (1999) Effects of temperature on slow and fast inactivation of rat skeletal muscle Na(+) channels. Am J Physiol 277:C937–947.

Schneider MF, Chandler WK (1973) Voltage dependent charge movement of skeletal muscle: a possible step in excitation-contraction coupling. Nature 242:244–246.

Schneider MF, Simon BJ (1988) Inactivation of calcium release from the sarcoplasmic reticulum in frog skeletal muscle. J Physiol 405:727–745.

Shang W, Lu F, Sun T, Xu J, Li LL, Wang Y, Wang G, Chen L, Wang X, Cannell MB, Wang SQ, Cheng H (2014) Imaging Ca2+ nanosparks in heart with a new targeted biosensor. Circ Res 114:412–420.

Uwera F, Ammar T, McRae C, Hayward LJ, Renaud JM (2020) Lower Ca2+ enhances the K+-induced force depression in normal and HyperKPP mouse muscles. J Gen Physiol 152.

van Lunteren E, Pollarine J, Moyer M (2006) Inotropic effects of the K+ channel blocker 3,4-diaminopyridine: differential responses of rat soleus and extensor digitorum longus. IEEE Trans Neural Syst Rehabil Eng 14:419–426.

Wang ZM, Messi ML, Delbono O (1999) Patch-clamp recording of charge movement, Ca2+ current, and Ca2+ transients in adult skeletal muscle fibers. Biophys J 77:2709–2716.

Yensen C, Matar W, Renaud JM (2002) K+-induced twitch potentiation is not due to longer action potential. Am J Physiol Cell Physiol 283:C169–177.

